# Activation of peroxisome proliferator-activated receptor-γ prevents monocrotaline-induced pulmonary arterial hypertension by suppressing of nuclear factor-kappa B mediated autophagy

**DOI:** 10.1101/2023.04.11.536344

**Authors:** Peng Zhang, Congcong Shi, Juntao Song, Tongbao Dong, Kaixiao Zhang

## Abstract

To investigate the molecular mechanisms underlying autophagy inducing pulmonary vascular remodeling and rosiglitazone inhibiting pulmonary arterial hypertension (PAH). Monocrotaline (MCT) was intraperitoneally injection to induce the rat PAH model. The right ventricular hypertrophy index (RVHI), right ventricle systolic pressure (RVSP), percentage of medial wall thickness (%MT) and histomorphologic analyses were performed to evaluate the development of PAH. The translocation of nuclear factor-kappa B (NF-κB) p65 subunit from cytosol to nucleus, the protein expression levels of LC3B, Beclin 1 and RND3 were determined by western blot. Furthermore, NF-κB inhibitor, pyrrolidine dithiocarbamate (PDTC) and peroxisome proliferator-activated receptor-γ (PPARγ) activator, rosiglitazone, were used to inhibit the activation of NF-κB and activate PPARγ signaling, respectively. MCT injection dramatically induced PAH models in rats as manifested by the increased RVSP, RVHI, and %MT. In addition, the activation of NF-κB and autophagy were significantly enhanced and the RND3 were markedly decreased in MCT-induced PAH in rat. However, these effects could be significantly suppressed either by the supplementation of PDTC or rosiglitazone. NF-κB promotes the development of PAH by activation of autophagy and consequent down-regulation of RND3 expression. Activation of PPARγ suppresses autophagy by inhibiting NF-κB in MCT-induced PAH. Our results not only uncovered the mechanisms of PPARγ activator in the protection of PAH but also provided potential therapeutic target for PAH treatment.

## INTRODUCTION

Pulmonary arterial hypertension (PAH), a rare but life-threatening dyspnoea-fatigue syndrome, which is characterized by a sustained and uncontrolled rise in pulmonary artery resistance and pulmonary vascular pressure, resulting in rapid progression of right ventricular hypertrophy, right ventricular systolic blood pressure, and eventually, heart failure and death [1, 2]. At present, the pathological mechanisms of PAH are remain largely unknown, however, genetic and epigenetic modifications, inflammatory and oxidative stress, pulmonary arterial endothelial cells dysfunction, metabolic and hormone imbalance, are considered as the mean risk factors [3]. In addition, it was believed that pulmonary vascular remodeling, the most prominent pathological feature of PAH, could be attributed to the proliferation of pulmonary arterial smooth muscle cells (PASMCs) [4]. Considering that the incidence of PHA is increasing rapidly in the world and have no efficient cure methods and may impose economic burden both on patient itself and healthcare system [5], therefore, elucidating the molecular mechanisms underlying the PASMCs proliferation and pursuing appropriate targets to inhibit vascular remodeling are of great meanings for the treatment of PAH.

Autophagy is a cellular self-degrading system, which is primordially conserved in eukaryotic, controls the degradation of the redundant or the damaged cellular components for renewal [6]. Autophagy is necessary to maintaining cellular homeostasis and energetic balance under stressful conditions and plays an important role in cell survival and maintenance [6, 7]. Autophagy dysfunction is highly associated with major human diseases such as cancer, neurodegenerative diseases, cardiomyopathy, diabetes, liver disease, autoimmune diseases, and infections [8].

Recent studies have suggested that activation of autophagy could induce the development of PAH in rats, and inhibition of autophagy decreases the proliferation of PASMCs in vitro [9-11]. However, the molecular mechanisms underlying autophagy in PAH are still largely unknown.

Nuclear factor-kappa B (NF-κB), is a conserved multifunctional transcription factor, plays a vital coordinating role in innate and adaptive immune responses, and has been implicating in a broad range of biological/pathological processes, including inflammation, cell proliferation, differentiation, apoptosis and cancers [12-14]. Previous studies has discovered that NF-κB promoted hypoxia-induced PAH by increasing the expression of dipeptidyl peptidase-4 (DPP4) in PASMCs and increased angiogenesis and pulmonary cell proliferation in PAH patients [15, 16]. Furthermore, NF-κB has been reported to induce autophagy activation in hyperlipidemia-induced cardiac remodeling [17]. However, whether NF-κB mediates the activation of autophagy in MCT-induced PAH is still unknown.

Peroxisome proliferator-activated receptor-γ (PPARγ), a ligand activated transcription factor that can be activated by peroxisome proliferators, plays key roles in multiple diseases including obesity, diabetes, cancer, and inflammatory diseases [18-20]. It has been demonstrated that PPARγ initiates a vascular regenerative program to reverse PAH, and prevents right heart failure via fatty acid oxidation [21]. Meanwhile, PPARγ has been demonstrated to interact with multiple signaling pathways, including B-cell lymphoma-2 (BCL2), NF-κB, p53, and Cyclin D1[22]. In addition, a study has reported that activation of PPARγ inhibits autophagy in myocardial hypertrophy [23]. However, it is unclear whether PPARγ modulates the activation of autophagy, thus the proliferation of PASMCs and pulmonary vascular remodeling to confer its overall beneficial effects on PAH.

In this study, we used rat models of MCT-induced PAH to examine the above issues and to gain more insight into the role of autophagy in the course of disease of PAH and, hopefully, to understand the underlying molecular mechanisms.

## MATERIALS AND METHODS

### Animals

Healthy male Sprague-Dawley (SD) rats (n = 50, 180-220g) with similar baseline characteristics were randomly assigned into 5 groups, namely, Control group, MCT group, MCT and PDTC group, MCT and HCQ group, MCT and rosiglitazone treatment group. All the rats were fed at 22 ± 2°C for 12 hours with light and dark cycles and free access to food pellets and water. All animal experiments were conducted in the Experimental Animal Center of Zibo 148 Hospital, in strict accordance with the "Guide for Experimental Animal Care and Use" of the Center. All protocols used in this study have been approved by the Experimental Animal Management Committee of our Hospital.

### Generation of PAH models and drug administration

MCT (Sigma-Aldrich, USA) was dissolved into 0.1 mol/L HCl, then adjusted the pH to 7.4 using 0.1 mol/L NaOH, and the final MCT concentration was 30 mg/mL. HCQ Sulfate tablets was the product of Shanghai Pharmaceutical Co., LTD, PDTC was purchased from Sigma-Aldrich, and Rosiglitazone tablets was purchased from Chengdu Hengri Pharmaceutical Co., LTD. These medicines were dissolved in 0.9% sodium chloride at final concentrations of 20 mg/mL, 50 mg/mL and 2.5 mg/mL, respectively. After one week of adaptive feeding, 60 mg/kg MCT was injected intraperitoneally to the rats for once to induce PAH models. After MCT injection, rats in the PDTC intervention group were intraperitoneally injected with 100mg/kg PDTC every day for a total of 28 days. HCQ and Rosiglitazone intervention groups were orally given 40mg/kg of HCQ and 5mg/kg of rosiglitazone per day for a total of 28 days, respectively. Control rats were received oral force-feeding or intraperitoneal injection of 0.9%NaCl throughout the experiment.

### Measurement of RVSP and RVH

On day 28, all surviving animals were anesthetized with isoflurane inhalation. After stable anesthesia, a polyethylene catheter was inserted through the isolated right internal jugular vein into the right ventricle (RV). The simulated signal of the right ventricular systolic blood pressure (RVSP) was detected by a Grass lie detector. After hemodynamic determination, the heart was extracted and dissected and the RV, left ventricle (LV), and interventricular septum (S) were dissected and weighed, respectively. The ratio of RV weight to LV plus S [RV / (LV+S)] was calculated to evaluate the index of right ventricular hypertrophy (RVHI).

### Histologic analysis

Marginal right lower pulmonary lobes from each rat were immersed immediately into 10% paraformaldehyde for fixation and then embedded with paraffin. The fixed pulmonary tissues were cut into 5 μm thickness and then dewaxed by adding xylene, anhydrous ethanol, 95% alcohol, 90% alcohol, 80% alcohol, 70% alcohol, and distilled water. Slices were dyed with Harris hematoxylin for 3-8min, washed with tap water, differentiated with 1% hydrochloric acid alcohol for several seconds, rinsed with tap water, returned blue with 0.6% ammonia water, and rinsed with running water. Slices were putted into eosin dye solution and stain for 1-3min. The slices were sequentially putted into 95% alcohol, anhydrous ethanol and xylene dehydration for transparent. After slightly dry and neutral gum seal. The slices were ready for microscopic examination, image acquisition. The percentage medial wall thickness (%MT) of vessels (20–70 μm diameters) was calculated by the method as follows: %MT = (2 × medial wall thickness) ×100/external diameter.

### Western blot analysis

Western blot was performed according the previous report [24]. Shortly, rat lung tissue was thoroughly lysed with RIPA after tissue homogenization. Nuclear proteins were extracted using nuclear protein extraction kits. The concentrations of total protein were quantified by BCA method. The proteins were loaded and separated by PAGE protein gel, and then transferred to polyvinylidene fluoride (PVDF) membrane. The membrane was blocked with 5% skim milk, and the primary antibody was incubated at 4[. After the secondary antibody was combined, the protein bands were detected by chemiluminescence method.

### Statistical analysis

All the data are expressed as mean ± standard deviation (SD), and are representative of three independent experiments. Multiple groups comparison using One-way ANOVA. P < 0.05 was considered differ significantly.

## RESULTS

### Activation of PPARγ prevented the increase of MCT-induced RVSP and RVHI

The RVSP values of rats in each group were detected by right heart catheterization. As shown in **Fig. 1A**, the RVSP was increased to 45.30 ± 2.11 mmHg in MCT-treated rats, which was significantly higher than that in control rats (22.62 ± 1.31 mmHg) (P<0.05; P < 0.05 versus control; **Fig. 1B**), suggesting that PAH was successfully induced in rats. Whereas, administration of PAH rats with PDTC or HCQ dramatically reduced RVSP values to 32.02 ± 1.98 mmHg and 35.21 ± 1.51 mmHg (P<0.05 vs. MCT-treated group; **Fig. 1B**), respectively. Furthermore, administration of pioglitazone in MCT-treated rats significantly reduced RVSP values to 30.06 ± 1.71 mmHg (P<0.05 vs. MCT-treated group; Fig.1B).

**Fig. 1.**
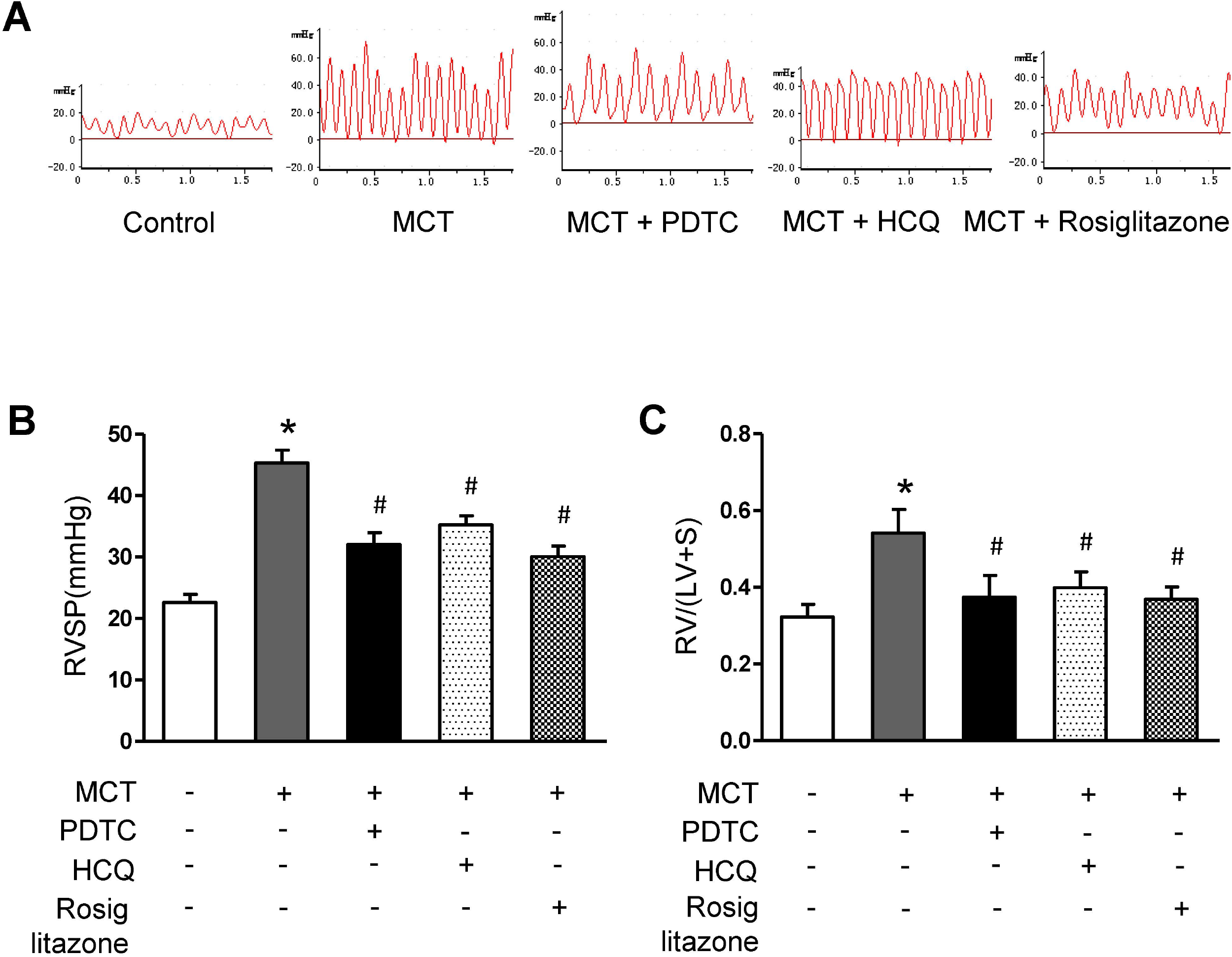
Activation of PPARγ prevented the increase of MCT-induced RVSP and RVHI. (A) The RVSP in different groups of rats was detected by right heart catheterization. (B) Change of RVSP in different groups of rats (n = 10-15). (C) Change of RVHI in different groups of rats (n = 10-15). \**P* < 0.05 versus control group; ^#^*P* < 0.05 versus MCT group.

Similar results were observed in RVHI. In MCT-treated PAH rats, RVHI was dramatically increased to 0.54 ± 0.06 versus 0.32 ± 0.03 in control group (P < 0.05 versus control; **Fig. 1C**). However, treatment of PAH rats with PDTC or HCQ significantly decreased RVHI values to 0.37 ± 0.06 and 0.40 ± 0.04 (P<0.05 vs. MCT-treated group; **Fig. 1C**), respectively. In conclusion, these results indicated that inhibition of NF-κB or autophagy process could effectively protected the development of PAH in rats. In MCT and pioglitazone-treated rats, the RVHI was declined to 0.37 ± 0.03 (P<0.05 vs. MCT-treated group; **Fig. 1C**), suggesting that activation of PPARγ by pioglitazone could significantly prevented the progression of PAH in rats.

### Activation of PPARγ prevented the increase of MCT-induced pulmonary arterial remodeling

As shown in **Fig. 2A and B**, medial wall thickness in small pulmonary arteries in MCT-treated rats [(36.21 ± 4.20) %] was markedly elevated when compared with that of control rats [(24.48 ± 2.64) %] (P < 0.05 versus control), this was accompanied with the increased number of PASMCs in the medial layer of the small pulmonary artery (**Fig. 2B**). However, after the usage of PDTC or HCQ, medial wall thickness dramatically decreased to (29.21 ± 4.00) % and (30.01 ± 3.21) %, respectively (P<0.05 vs. MCT-treated group; **Fig. 2A, B**). PDTC or HCQ also prevented MCT-induced the increase of PASMCs number. These results indicated that inhibition of NF-κB or autophagy could significantly suppressed the pulmonary arterial remodeling in MCT-induced PAH. What’s more, treatment of PAH rats with pioglitazone dramatically decreased medial wall thickness to (27.98 ± 3.01) % (P < 0.05 versus MCT-treated rats; **Fig. 2A, B**), this was accompanied with the decrease of PASMCs number. In conclusion, these results indicated that activation of PPARγ could significantly suppressed MCT-induced pulmonary arterial remodeling in rats.

**Fig. 2.**
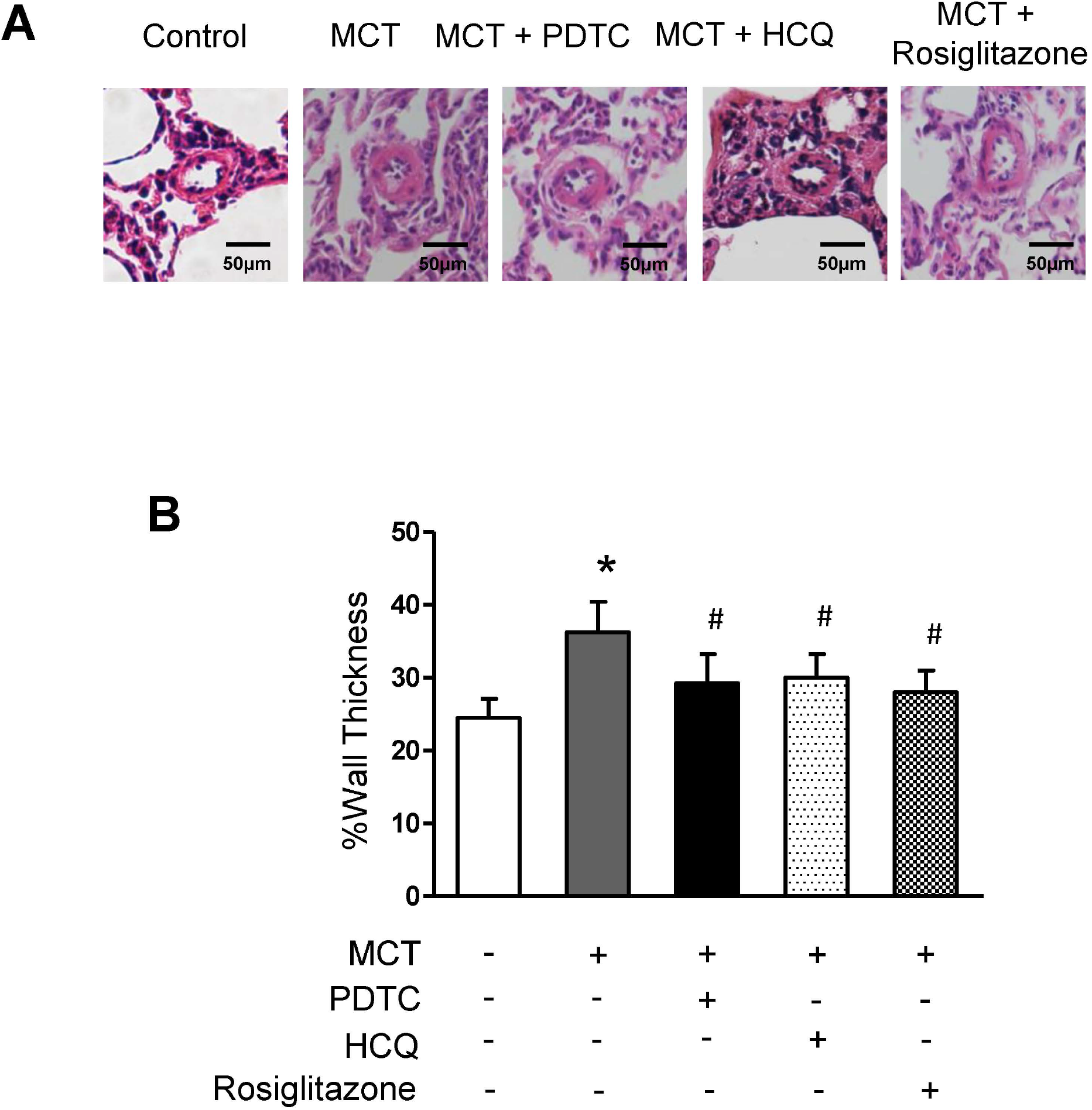
Activation of PPARγ prevented the increase of MCT-induced pulmonary arterial remodeling. (A) Hematoxylin and eosin staining of small pulmonary arteries in different groups (n = 10-15) (magnification × 400). (B) Quantitative analysis of the medial wall thickness of pulmonary arteries (n = 10-15). \**P* < 0.05 versus control group; ^#^*P* < 0.05 versus MCT group.

### Activation of PPAR**γ** inhibited NF-**κ**B activation in MCT-induced PAH model

In order to clarify whether NF-κB is activated in MCT-induced PAH rats, the translocation of NF-κB p65 subunit from cytosol to nucleus was measured using western blotting. As shown in **Fig. 3**, the protein expression level of NF-κB p65 subunit in the nuclear was increased to 3.86-fold over control in MCT-treated rats (P < 0.05 versus control). Whereas, administration of PDTC, HCQ or pioglitazone significantly reduced nuclear NF-κB p65 subunit level in MCT-treated rats to 2.50-fold, 1.35-fold and 1.53-fold over controls, respectively, (P < 0.05 versus MCT-treated rats; **Fig. 3**). At the same time, the protein expression level of NF-κB p65 subunit in the cytosol was declined to 0.29-fold over controls in MCT-treated rats (P < 0.05 versus control). While treatment of PAH rats with PDTC, HCQ and pioglitazone increased cytoplasmic NF-κB p65 subunit level to 0.53-fold, 0.87-fold or 0.94-fold over control, respectively, (P < 0.05 versus MCT-treated rats; **Fig. 3**). These results indicated that activation of PPARγ could significantly suppressed NF-κB activation in MCT-induced PAH in rats.

**Fig. 3.**
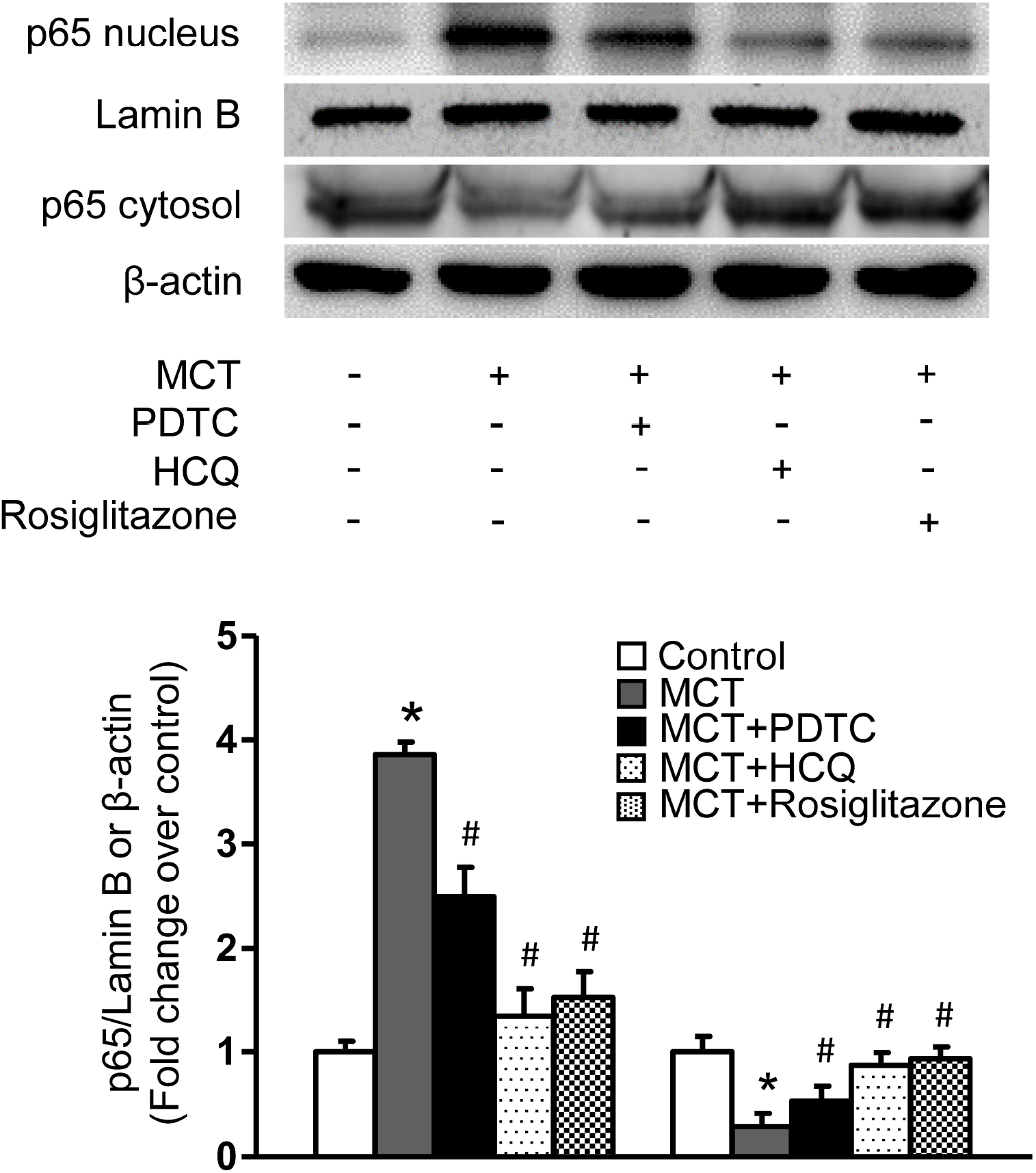
Activation of PPARγ inhibited NF-κB activation in MCT-induced PAH model. The protein level of the NF-κB p65 subunit in the nuclear and cytoplasmic fraction was determined using western blot and Lamin B and β-actin served as loading control for the nuclear and cytoplasmic protein, respectively (n=4 each group). \**P* < 0.05 versus control group; ^#^*P* < 0.05 versus MCT group.

### Activation of PPARγ inhibited autophagy activation in MCT-induced PAH model

To determine whether autophagy is activated in MCT-induced PAH rats, the protein levels of LC3B and Beclin 1 were measured using western blotting (**Fig. 4**). In MCT-treated rats, the protein expression levels of LC3B and Beclin 1 were increased to 2.27-fold and 2.07-fold over controls, respectively (P < 0.05 versus control; **Fig. 4A and B**). Whereas, administration of PDTC, an inhibitor of NF-κB, could significantly decreased the protein expression levels of LC3B and Beclin 1 to 0.47-fold and 1.07-fold over controls (P < 0.05 versus MCT-treated rats; **Fig. 4A and B**), respectively.

**Fig. 4.**
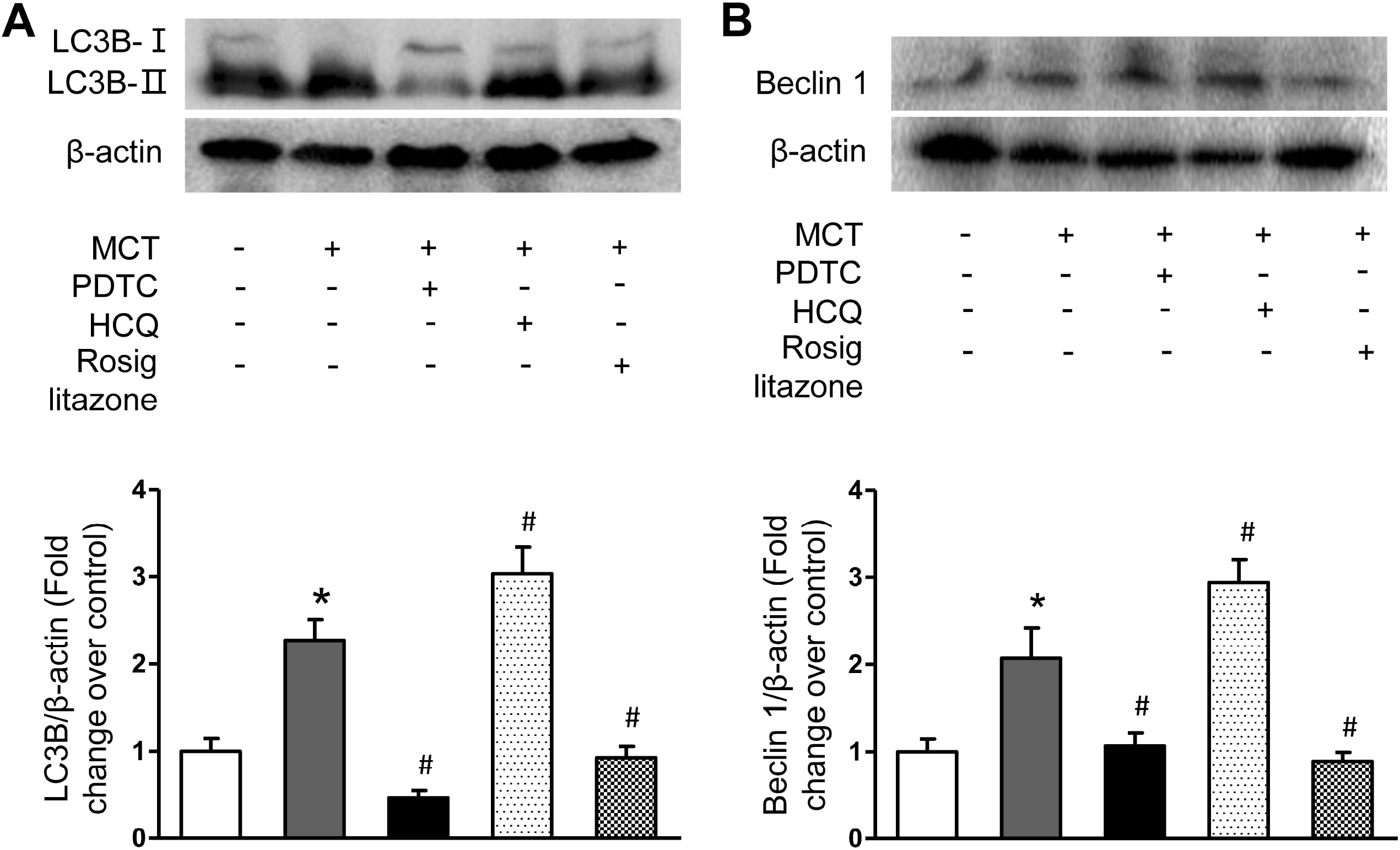
Activation of PPARγ inhibited autophagy activation in MCT-induced PAH model. The protein levels of LC3B (A) and Beclin 1 (B) in lung tissues from different groups were determined using immunoblotting. Representative western blot and quantification of bands are shown. (n = 4 each group). \**P* < 0.05 versus control group; ^#^*P* < 0.05 versus MCT group.

HCQ, an autophagy inhibitor, which protects autophagy by inhibiting the acidification of the lysosomes that fuse with the autophagosomes, thereby preventing the degradation of metabolic stress products and enhancing the accumulations of autophagy markers, such as LC3A, LC3B and Beclin 1 [25]. Treatment of PAH rats with HCQ further increased the protein expression levels of LC3B and Beclin 1 to 3.03-fold and 2.94-fold over control (P < 0.05 versus MCT-treated rats; **Fig. 4A and B**). These results implied that the increased autophagy activity could attributed to the aberrant activation of NF-κB signaling in MCT-induced PAH rats. In addition, the administration of pioglitazone could significantly decreased the protein expression levels of LC3B and Beclin 1 to 0.93-fold and 0.89-fold over controls (P < 0.05 versus MCT-treated rats; **Fig. 4A and B**), suggesting that activation of PPARγ inhibited the activation of autophagy in MCT-induced PAH.

Meanwhile, immunohistochemistry was used to detect the protein expression of LC3B. The percentage of LC3B-positive PASMCs was increased from (4.1 ± 1.2) % in control rats to (20.2 ± 2.0) % in MCT-treated PAH rats (P < 0.05 versus control) (**Fig.5**). Whereas treatment of PAH rats with PDTC significantly decreased the percentage of LC3B-positive PASMCs to (13.6 ± 2.1) % (P < 0.05 versus MCT-treated rats, **Fig. 5**). Furthermore, administration of HCQ dramatically increased the ratio of LC3B-positive PASMCs to (27.1 ± 2.6) % (P < 0.05 versus MCT-treated rats, **Fig. 5**), whereas the percentage of LC3B-positive PASMCs was decreased to (14.5 ± 2.2) % in MCT and pioglitazone co-treated rats (P < 0.05, **Fig. 5**). In conclusion, these results suggested that activation of PPARγ could significantly inhibited autophagy by suppressing of NF-κB nuclear translocation in MCT-induced PAH in rats.

**Fig. 5.**
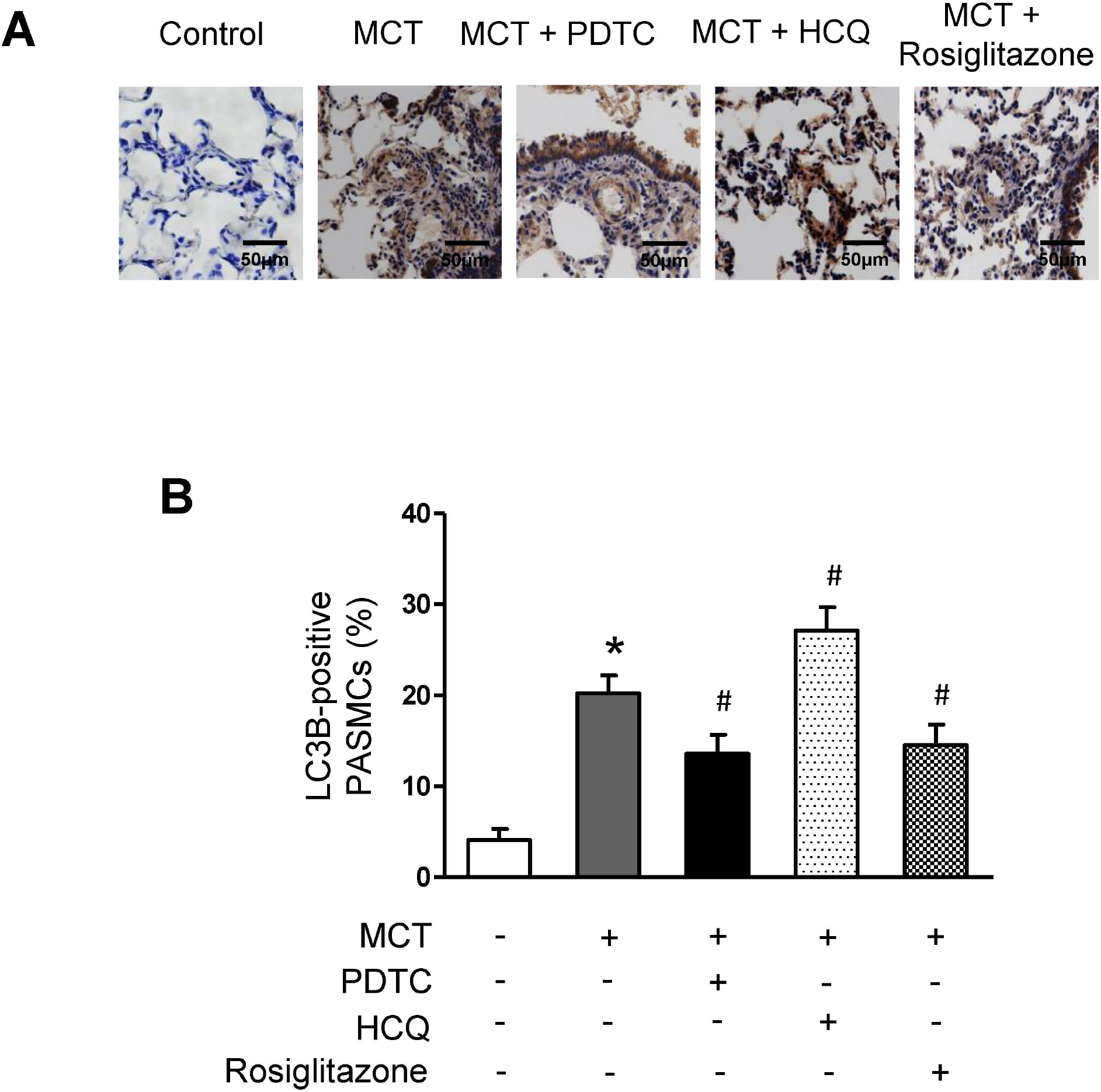
Activation of PPARγ inhibited the percentage of LC3B-positive PASMCs in MCT-induced PAH model. (A) Immunohistochemistry was used to detect the protein level of LC3B in the lung tissues from different groups; cells with brown-stained cytoplasm in the medial layer of pulmonary artery are LC3B-positive PASMCs. (B) The percentage of LC3B-positive PASMCs in different groups (25 arteries each slide, diameter 20-100 μm). \**P* < 0.05 versus control group; ^#^*P* < 0.05 versus MCT group.

### Activation of PPARγ restored the expression of RND3 in MCT-induced PAH model

RND3, an atypical member of Rho family, which has been shown to regulate a variety of biological activities, such as cell proliferation, migration, and apoptosis [26, 27]. Therefore, we examined the expression of RND3 in lung tissue of PAH rat. As shown in **Fig. 6**, the protein expression level of RND3 was significantly declined to 0.26-fold over controls in MCT-induced PAH rats (P < 0.05 versus control), while administration of PAH rats with either PDTC or HCQ could significantly increased the protein expression level of RND3 to 0.90-fold or 0.92-fold over control (P < 0.05 versus MCT-treated rats). These results implied that inhibition of NF-κB or autophagy could abrogated the reduction of RND3 in MCT-induced PAH rats. In addition, RND3 protein level was also increased to 0.906-fold in MCT and pioglitazone treated PAH rats (P < 0.05 versus MCT-treated rats; **Fig. 6**), suggesting that activation of PPARγ could restored the reduction of RND3 in MCT-induced PAH.

**Fig. 6.**
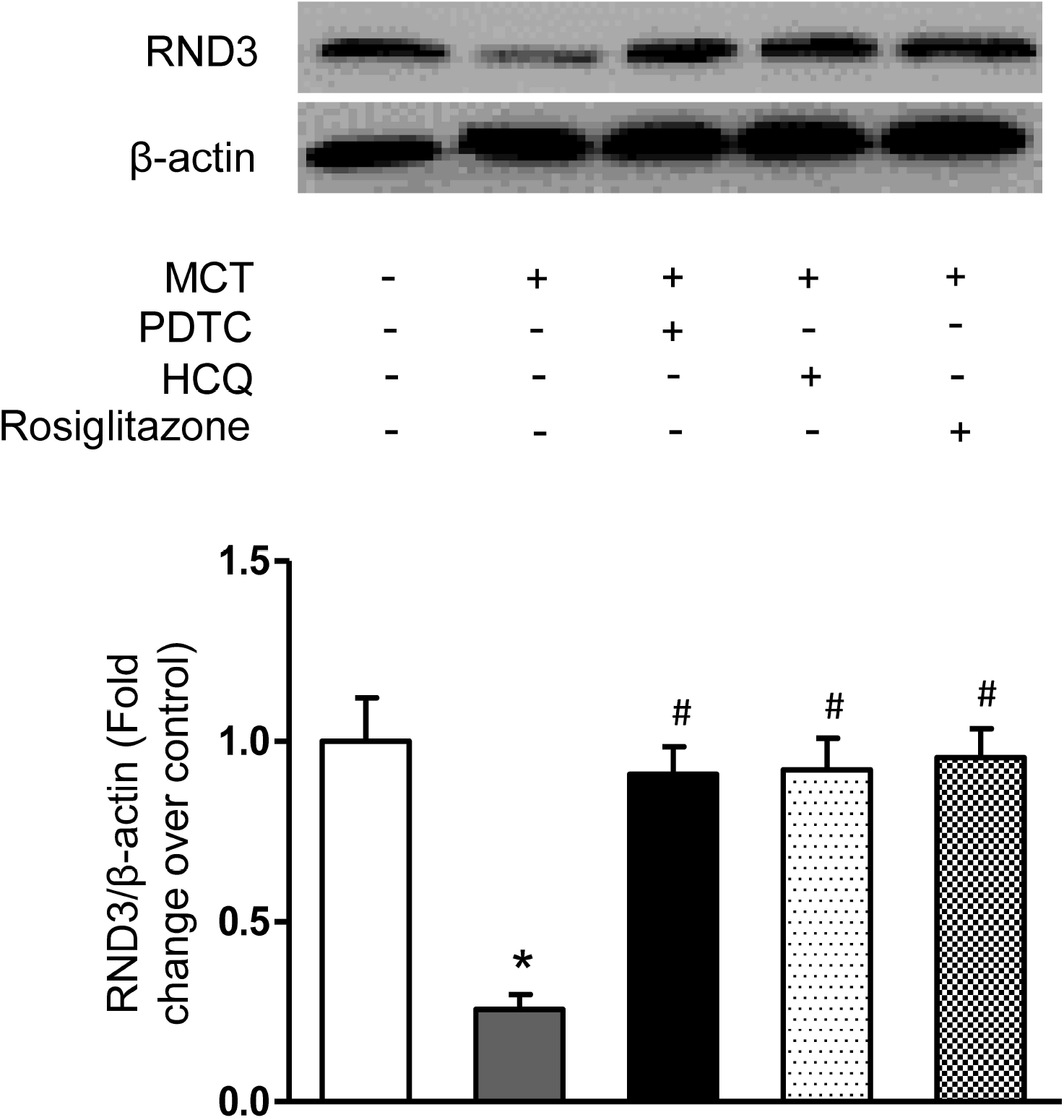
Activation of PPARγ abrogated the expression of RND3 in MCT-induced PAH model. The representative western blot of RND3 protein level in lung tissues from different groups and its quantification analysis. β-actin served as a loading control (n=6 each group). \**P* < 0.05 versus control group; ^#^*P* < 0.05 versus MCT group.

## DISCUSSION

In this work, we discovered that MCT injection could dramatically induced PAH-like models in rats, which manifested by the increased levels of RVSP, RVHI, and %MT. What’s more, the activation of NF-κB and autophagy were significantly enhanced and the RND3 were markedly decreased in MCT-induced PAH in rats. However, these effects could be significantly suppressed either by the supplementation of PDTC or rosiglitazone. Our results not only revealed the mechanisms of the interplay of PPARγ/NF-κB/autophagy axis in the regulating of PAH, but also provide potential therapeutic targets for the prevention and treatment of PAH.

Autophagy, a major degradation pathway in eukaryotic cells, which not only transports abnormal substances from cytoplasm to lysosomes for degradation but also plays an important role in the process of health and disease [28, 29]. Under normal circumstances, cell autophagy are rarely occur in cells, only if there is a predisposing factors, which including nutrient starvation, hypoxia, protein aggregates, damaged organelles, or, most commonly, intracellular pathogens infection [30, 31]. Previously, it has been already shown that autophagy could induces the proliferation of PASMCs and pulmonary artery remodeling in PAH animals [32, 33]. Whereas, inhibition of autophagy could suppressed sphingosine-1-phosphate (S1P)-stimulated PASMCs proliferation and MCT-induced pulmonary artery remodeling [34-36]. Furthermore, it has been reported that activation of autophagy strengthened MCT-induced PAH might through the regulation of FOXM1-FAK signaling pathway as researchers have discovered that both FOXM1 and FAK inhibition not only suppressed MCT-induced autophagy activation, but also decreased RVSP, RVHI and %MT [37]. In this study, we broadened these notions by demonstrating that the protein levels of LC3B and Beclin 1 were increased in MCT-induced PAH rats, and inhibition of autophagy by HCQ suppressed pulmonary artery remodeling and the development of PAH.

NF-κB, a protein complex, is stimulated by a variety of stimuli, such as hypoxia, inflammatory cytokines, or intracellular pathogens infection [38]. When NF-κB is activated, the two subunits of NF-κB, NF-κB/IκB complex will separate, and this will result in the release of p65/p50 heterodimers subunits. Furthermore, p65/p50 heterodimers is transferred from cytoplasm to the nucleus to regulate multiple biological functions, including immune response, inflammation, and cell survival [39, 40]. One of the characteristics of PAH is the remodeling of pulmonary artery extracellular matrix (ECM) [41]. ECM remodeling and increased pulmonary artery stiffness mechanically activate various signaling pathways, such as transforming growth factor-β (TGF-β), and NF-κB [42]. Therefore, inhibition of ECM remodeling and mechanical conduction is expected to prevent and even reverse experimental PAH [41]. Previously, Hong and his colleagues used single-cell transcriptomics discovered that hypoxic non-classical monocytes and MCT conventional dendritic cells (DC) showed extremely strong activation of the NF-κB pathway [43]. Furthermore, studies in NF-κB was found to be activated in endothelin-1-induced proliferation of PASMCs and MCT-induced PAH in mice [44, 45]. In our study, we found that NF-κB activity is up-regulated in MCT-induced PAH, which was accompanied with increased pulmonary vascular remodeling. The role of NF-κB in the activation of autophagy is conflicting. A previous report has shown that NF-κB induces the activation of autophagy in pancreatic ductal adenocarcinoma cells [46]. However, other studies have discovered that NF-κB inhibits autophagy in several cancer cell lines [47]. In our study, we found that NF-κB inhibitor PDTC reduced autophagy-related marker proteins (LC3B and Beclin 1) and alleviated pulmonary artery remodeling.

Peroxisome proliferator-activated receptors (PPARs), is a ligand-activated transcription factors, which is involved in the regulating of glucose and lipid homeostasis, inflammatory response, proliferation and differentiation [48, 49]. PPARγ agonist pioglitazone could reversed pulmonary vascular pressure and prevented right heart failure by facilitating fatty acid oxidation [50]. Importantly, a previous study has suggested that myostatin could inhibited NF-κB signaling pathway by promoting the expression of PPARγ, and then suppressed excessive autophagy in cardiac hypertrophy [23]. In this study, we have not only confirmed that activation of PPARγ could inhibited the translocation of NF-κB p65 subunit from the cytosol to the nucleus, and then reduced autophagy-related marker proteins in MCT-induced PAH but have also verified that NF-κB inhibition could dramatically decreased the symptoms of experimental PHA in rats. These results imply that activation of PPARγ could inhibited the activation of autophagy via the suppression of NF-κB activation and translocated to the nuclear in MCT-induced PAH.

RND3, an anti-proliferative protein, regulates multiple cell physiological functions, which including cell proliferation, growth, apoptosis, and differentiation [26, 51]. On account of the capabilities of RND3 to inhibit the proliferation of cells, thus, RND3 is usually significantly down-regulated in a variety of human cancer cell lines [52, 53]. In this study, we indicated that the protein level of RND3 was significantly decreased in MCT-induced PAH. In addition, autophagy has been shown to promote the degradation of RND3 by the autophagy-lysosomal pathway in gastric cancer cells [54]. Our studies were in line with the previous results as we verified that HCQ could restored MCT-induced down-regulation of RND3 expression. These results suggest that inhibition of autophagy restores the protein level of RND3 in MCT-induced PAH.

## CONCLUSION

We identify that NF-κB induces the development of PAH by stimulating autophagy and subsequent downregulation of RND3 expression. Activation of PPARγ suppresses pulmonary artery remodeling by inhibiting NF-κB and thus autophagy. Our study may contribute to the development of more effective PAH therapeutic strategies.

### Contributions

All authors contributed to the study conception and experiment design. Material preparation, data collection and analysis were performed by Peng Zhang, Congcong Shi, Taojun Song, and Tongbao Dong. The first draft of the manuscript was written by Peng Zhang, Kaixiao Zhang and all authors commented on previous versions of the manuscript. All authors read and approved the final manuscript.

### Ethics declarations

The authors have no relevant financial or non-financial interests to declare. All procedures performed in studies on animals were in compliance with ethical standards of the institution in which the studies were conducted and with the approved legal acts of the Russian Federation and international organizations.

## Abbreviations

BCL2: B-cell lymphoma-2
DPP4: dipeptidyl peptidase-4
ECM: extracellular matrix
LV: left ventricle
MCT: Monocrotaline
NF-κB: nuclear factor-kappa B
PAH: pulmonary arterial hypertension
PASMCs: pulmonary arterial smooth muscle cells
PDTC: pyrrolidine dithiocarbamate
PPARγ: peroxisome proliferator-activated receptor-γ
PVDF: polyvinylidene fluoride
RND3: Rho family GTPase 3.
RV: right ventricle
RVHI: right ventricular hypertrophy index
RVSP: right ventricle systolic pressure
%MT: percentage of medial wall thickness
SD: standard deviation
TGF-β: transforming growth factor-β.

